# Modulation of Purkinje Cell Inhibition by Stem Cell Factor

**DOI:** 10.1101/2024.12.09.627581

**Authors:** Tariq Zaman, Jessica Lahr, Suraj Cherian, Lucas Pozzo-Miller, Michael R Williams

## Abstract

Target derived factors influence the specification, maintenance, and modulation of synaptic connectivity. The transmembrane protein, Kit Ligand, and Kit receptor tyrosine kinase are differentially expressed in connected neurons. In development and postnatal periods, these proteins maintain connectivity between cerebellar Purkinje cells (PC) that express Kit Ligand, and presynaptic Molecular Layer Interneurons (MLI) expressing Kit. In this study, it is demonstrated that Stem Cell Factor (SCF), the active extracellular domain of Kit Ligand, produces a potent potentiation of inhibition upon Purkinje Cells. The SCF enhancement of inhibition required presynaptic Kit, produced long term suppression of PC firing, and was associated with a specific potentiation of basket cells of the MLI1 subtype. It is posited that SCF exerts a postsynaptic effect involving enhanced sensitivity of somatic PC GABA_A_ receptors. This work demonstrates that the SCF/Kit axis modulates synaptic function in adult tissue.

## Introduction

The receptor tyrosine kinase, Kit, is expressed in discrete populations of neurons across a variety of circuits and species. Mutation, inhibition, or knockout of Kit can produce neurodevelopmental phenotypes, suggesting that the activation of Kit kinase regulates neuronal development or physiology. The endogenous agonist of Kit is Kit Ligand, a single pass transmembrane protein that exists in at least two isoforms differing in their inclusion of an exon encoding a sheddase/cleavage motif that facilitates the liberation of soluble Kit Ligand, also known as, and referred herein to as, Stem Cell Factor (SCF)(1, 2). As Kit Ligand and Kit receptor protein expression are maintained throughout maturity, it is feasible that the degree of SCF mediated Kit activation can regulate the function of mature neural circuits; given the broad role of receptor kinases in neuromodulation.

Within the brain of humans and multiple model organisms, Kit Ligand is highly expressed by cerebellar Purkinje Cells (PCs), and Kit receptor is highly expressed by cerebellar cortex molecular layer interneurons (MLIs) that provide GABAergic inhibition to PCs. We previously established through conditional knockout mice that MLI mediated PC expression was impaired by the developmental knockout of Kit Ligand or of Kit. Furthermore, we found that delaying PC Kit Ligand knockout until adulthood, also resulted in a decrease in synaptic inhibition of PCs. Although PCs receive inhibition from multiple cell types, the principal source of their GABAergic input is from MLI Type 1s (MLI1), which synapse upon each other to promote MLI1 synchrony, and upon PCs to modulate their firing (rate and variability) and to gate their plasticity. The degree of MLI mediated PC inhibition can be modulated by endogenous cascades including glutamate spill over, cannabinoids, and PC derived peptides. Here we sought to determine if SCF mediated Kit activation may be a mechanism for MLI1 mediated PC inhibition.

## Methods

### Animals

All procedures performed at Michigan State University were approved by the Institutional Animal Care and Use Committees of Michigan State University, which is accredited by the Association for Assessment and Accreditation of Laboratory Animal Care. Mice were on a 12-hour dark/light cycle, with ad libitum food and water. Wild type C57BL/6J male and female mice were used for most experiments. Kit receptor floxed homozygous mice (Kit^tm1c(EUCOMM)Mirow^/JN, PMID:(3)), some carrying a Pax2 transgene (Tg(Pax2-Cre)1Akg/Mmnc, RRID:MMRRC_010569-UNC (4)) were used for conditional cerebellar knockout of Kit as previously described. Animals were genotyped in-house using standard PCR-based methods, or via Transnetyx.

### Acute slice preparation

Avertin-anesthetized mice were perfused with a carbogen-equilibrated ice-cold slicing solution containing (in mM) 234 sucrose, 10 MgS0_4_, 2.5 KCl, 1.25 NaH_2_PO_4_, 24 NaHCO3, 0.5 CaCl_2_, and 11 glucose (5). In the same solution, using a vibrating tissue slicer (Leica VT1200S), we generated 300-micron-thick parasagittal slices of the cerebellum from animals at 26.1 ± 0.3, 49.9 ± 0.5 and 28.6 ± 0.3 days for MLIs, PCs and MLI-PC paired recording, respectively. Slices recovered at 34°C for 30 minutes in artificial cerebrospinal fluid (ACSF, described below) and then held at room temperature in the same ACSF for at least 30 minutes before recording.

### Extracellular Solutions

For single cell recordings, slices were incubated in carbogenated ACSF containing (in mM) 125 NaCl, 25 NaHCO_3_, 1.25 NaH_2_PO_4_, 2.5 KCl, 1 MgCl_2_, 1 CaCl_2_, and 25 glucose; recordings were performed in ACSF (32.7 ± 0.1°C) containing (in mM) 125 NaCl, 25 NaHCO_3_, 1.25 NaH_2_PO4, 2.5 KCl, 1 MgCl_2_, 2 CaCl_2_, and 25 glucose as described previously (3).For dual cell recordings, slices were incubated in the ACSF containing (in mM) 119 NaCl, 26.2 NaHCO_3_, 1 NaH_2_PO4, 2.5 KCl, 1.3 MgCl_2_, 1 CaCl_2_, and 11 glucose; recordings were performed in ACSF (32.7 ± 0.1°C) containing (in mM) 119 NaCl, 26.2 NaHCO_3_, 1 NaH_2_PO_4_, 2.5 KCl, 1.3 MgCl_2_, 2 CaCl_2_, and 11 glucose as described by (6).

### Patch-Clamp Conditions

MLIs and PCs of folia IV/V were targeted for recording by IR-DIC detected via a SciCam Pro on a SliceScope Pro 6000-based rig (Scientifica). Recording electrodes were pulled (Narishige, PC-100) from standard-wall borosilicate glass capillary tubing (G150F-4, Warner Instruments) and had 4.5 to 5.5 and 2.5 to 3.0 MΩ tip resistance for MLIs and PCs, respectively. Inhibitory post synaptic currents (sIPSC and mIPSC) were recorded with an intracellular solution containing (in mM)140 CsCl, 4 NaCl, 0.5 CaCl2-2H2O, 10 HEPES, 5 EGTA, 2 Mg-ATP, and 0.4 Na-GTP, 2 QX-314. In whole-cell voltage-clamp mode, PCs were held at –70 mV; to isolate sIPSCs, NBQX (10 µM) and D-AP5 (50 µM) were added to the recording solution. To confirm GABA_A_-mediated sIPSCs, strychnine (1 µM) and CGP-52432 (2 µM) were further applied, and mIPSC recordings additionally included 1 µM tetrodotoxin (TTX). For action potential recordings, the intracellular solution contained (in mM) 140 K-gluconate, 10 KCl, 1 MgCl2, 10 HEPES, 0.02 EGTA, 3 Mg-ATP, and 0.5 Na-GTP as described previously (PMID:(5)). The internal pipette solution pH was adjusted to 7.35 with CsOH for IPSCs and mIPSCs, and with KOH for action potentials, while for all the osmolarity was adjusted to 300 mOsmolL-1 with sucrose. To examine SCF-mediated potentiation of synaptic inhibition, slices were treated with recombinant mouse SCF (150 ng/mL, PMID: 12137920); SCF was acquired as SRP3234-10UG, S9915-50UG, (MilliporeSigma) or 455-MC-500 (R&D Systems).

Spontaneous action potentials in PCs and from MLI from the lower third of the molecular were recorded in I = 0 mode. To examine SCF-potentiated synaptic inhibition, connected pairs of MLI and PC were held at -60mV, 300-ms depolarizing current was injected in PC, and 100 ms later a 200 HZ train of current pulses (2 ms) was delivered in the MLI. The firing frequency of PC before and during MLI firing was compared between Control and SCF condition. For each cell, 3-5 sweeps were recorded with 7 s inter-sweep interval. A square-wave voltage stimulation pulse was utilized to determine input resistance and cell capacitance. PCs with an access resistance of 10–20 (15.9 ± 0.5) MΩ or MLIs with an access resistance < 30 (23.6 ± 0.6) MΩ were considered for recording. Recordings with > 20% change in series resistance were excluded from analysis. Signals were acquired at 10 kHz with a low-noise data acquisition system (Digidata 1550B) and a Multiclamp700-A amplifier and were analyzed using pClamp11.1 (Molecular Devices) after low pass Bessel (eight-pole) filtration (3 dB cutoff, filter 1 kHz). The minimum amplitude threshold for detecting IPSCs was 15 pA and for mIPSCs the cutoff was 8 pA. For determining frequency and amplitude, individual events longer than 1 ms were included while overlapping events were manually rejected; also rejected from individual event analysis was any mIPSC for which amplitude, rise, and decay metrics were not all available.

### Software

Electrophysiological data was analyzed by pClamp11.1, statistical analyses were conducted via GraphPad Prism 9, and figures were created with BioRender, PowerPoint, and GraphPad Prism 9.

## Results

It has previously been demonstrated that 30-minute bath application of recombinant SCF at 150 ng/mL produces a long term potentiation of excitatory glutamatergic neurotransmission in the tri-synaptic circuit of the hippocampus mice, owing to the enrichment of Kit in CA subfields (7–10). We therefore investigated if this experimental strategy would also potentiate inhibition of PCs, owing to enrichment of Kit in MLIs. We performed whole cell patch clamp recordings of PCs from acutely generated cerebellar slices of young adult mice that had been incubated for 30 minutes in a control ACSF solution or in the same plus 150 ng/mL of SCF. In “wild type” C57Bl6/J mice, SCF significantly increased the frequency of sIPSC recorded in PCs (11.27±1.18 vs 15.46±1.56 Hz, n= 18 vs 18, two-tailed t-test p=0.0267). A similar effect was observed in Kit floxed homozygous mice on a C57BL6/J background (10.34±1.01 vs 15.45±1.88 Hz, n=10 vs10; two-tailed t test p =0.028). The bath SCF mediated increase in PC sIPSC frequency was not observed however in Kit floxed homozygous mice expressing Pax2 Cre, which we have previously demonstrated accomplishes developmental knockout of Kit protein from the cerebellum REF (5.29±0.63 vs 5.43±1.12 Hz, n=11 vs 11; two-tailed t test p=0.91). These data suggest that SCF acts through Kit to potentiate synaptic inhibition of Purkinje cells. However, in these same studies, we did not detect a significant increase in the per-PC mean sIPSC amplitude. We therefore considered the possibility that our initial 30-minute time point was in the waning phase of the SCF response, or that the population variability within each condition obscured the differences between the conditions. We therefore investigated, on a per cell basis, the acute effects of SCF in a before vs after design.

We performed whole cell patch clamp recordings of PC from acutely generated cerebellar slices of young adult mice before and after the application of SCF. We determined the mean sIPSC frequency and amplitude at baseline, and at each of five one-minute intervals after initiating the SCF application. For the baseline timepoint, data was obtained from 17 PCs in the Control condition (Kit Floxed homozygous Cre negative animals), and from 10 PCs in the Kit KO condition (Kit Floxed homozygous, Pax2 Cre positive). We tested for significant effects using mixed-effects model (REML), with a matched design strategy (each cell, across time), without assuming sphericity, an alpha of 0.05, with fixed effects being Time in SCF (p=0.0436), Kit Condition (Control vs KO, p=0.0317), and Time x Kit interaction (p=0.0854). Using a Holm-Sidak multiple comparisons test, we determined that PC sIPSC frequency was significantly increased vs baseline at minutes 4 (p=0.0146) and 5 (p=0.0334) after the start of SCF; for the Kit KO condition, SCF had no significant effect at any timepoint (p≥0.683). Having determined that SCF produced a rapid Kit mediated potentiation of sIPSC frequency, we evaluated sIPSC amplitude in the same fashion. The fiexed effects were significant, Time in SCF (p=0.0104), and Kit Condition (p=0.0373), as was their interaction (p<0.0001), and we determined that SCF had a significant effect at minutes 3 (p=0.015), 4 (p=0.0091), and 5 (p=0.0047) vs baseline . For the Kit KO condition, SCF had no significant effect at any timepoint (p=≥0.60) In a separate set of experiments, we determined that the acute application of SCF still significantly enhanced the frequency of sIPSCs in PCs in the additional presence of blockers of glycinergic (strychnine) and metabotropic GABA receptors (CGP-52432), and we confirmed that these SCF potentiated synaptic events were blocked by the additional application of picrotoxin. From a sample size of 10 PCs, data were analyzed by repeated-measures ANOVA, not assuming sphericity; we found a significant difference between groups (Control=17.23±2.61Hz, SCF=25.03±4.53 Hz; Picrotoxin=1.55±0.62 Hz; p=0.0001). By Holm-Šídák’s multiple comparisons test we determined that Control vs. SCF (P=0.0204), Control vs. Picrotoxin p=0.0007, and SCF vs. Picrotoxin p=0.0009.

From these experiments, we determined that SCF produced a significant potentiation of fast synaptic GABAergic inhibitory drive to PCs that depended upon the Kit receptor. MLI mediated inhibitory drive to PCs is known to modulate PC firing rate and variability; we determined if SCF also modulated PC firing. We performed whole-cell patch clamp on PCs from C5Bl6/J mice before, and at 5 minutes after the application of SCF. The application of SCF significantly decreased mean PC firing rate (56.82±5.58 Hz vs 38.66 ± 5.45 Hz (SCF), p=0.0011, n=13) , significantly increased the mean inter-spike interval (19.55±2.07 ms vs 32.91± 5.67 Hz (SCF), p=0.0046, n=13), and significantly increased the variability of the inter-spike interval (0.129± 0.023 vs 0.186 ± 0.1257 (SCF), p=0.0327, n=13). These effects were analyzed by paired test of the average value per PC, during the baseline vs post-SCF period; firing rate was analyzed by two-tailed t-test, while ISI and ISI CV2 was analyzed using Wilcoxon matched-pairs signed rank test. Having determined that the SCF mediated facilitation of inhibition also leads to changes in PC firing dynamics, we determined whether these effects persisted over time.

We determined PC average firing rate during a Baseline period, at the end of a 5-minute 150ng/mL SCF or Bovine Serum Albumin vehicle (BSA) “Drug” application, and then at each minute for ten minutes of an ACSF Washout period. We analyzed the data using a Two-way Repeated Measures ANOVA with a matched design strategy (each cell, over time), without assuming sphericity, an alpha of 0.05, with fixed effects of Time (p=0.105), Drug (p=0.154), and Time x Drug interaction ( p<0.0001). We conducted Holm-Sidak multiple-comparisons test and determined that, compared to Baseline, PC firing rate was significantly different at every timepoint for the SCF condition (p≤0.0105), but not for the BSA condition (p≥0.996 at each timepoint vs Baseline). We thus interpreted that a transient SCF application led to a persistent suppression of PC firing for at least 10 minutes after the beginning of the washout period, and since this effect was not observed in the BSA condition, was unlikely to be due to rundown of PC firing.

We investigated if an increase in MLI firing rate was mechanistically involved in SCF potentiated PC inhibition. We recorded the per-cell average spontaneous firing rate of MLIs before vs after SCF. MLIs are a heterogenous population; we analyzed those with membrane input resistance less than, or greater than, 500 Ω·cm^2^. Those having a lower input resistance have been classified as MLI1s, which synapse upon other MLI1s to promote synchrony upon PCs to promote inhibition, while those with higher input resistance have been classified as MLI2s, which principally inhibit MLI1s (11, 12). We analyzed the data using a two-way repeated measures (per-cell) design, assuming sphericity with an alpha of P=0.05. The MLI type (Rm) was a significant source of variation (P<0.0001), while SCF (P=0.658) , Cell (P=0.699), and Cell x SCF interaction (P=0.195), terms were not significant sources of variation. Holm-Sidak multiple comparisons test confirmed that SCF had no significant effect in the MLI1 (Rm<500, 26.07 ±4.53 vs 23.26±3.49 Hz, n=10, P=0.374) or MLI2 (Rm<500, 26.35±5.18 vs 27.76±4.54 Hz, n=9 P=0.545) groups. Since SCF only non-significantly *reduced* MLI1 firing rate, it is very unlikely that *increased* MLI1 firing rate contributes mechanistically to SCF potentiated PC inhibition.

Towards investigating pre- vs postsynaptic mechanisms, we recorded miniature inhibitory post-synaptic currents (mIPSC) from PC before vs after SCF. The average frequency of mIPSC was not changed (5.45±0.34 Control vs. 5.61±0.58 Hz SCF , two-tailed paired t-test p=0.78, n= 14 cells). However, the average amplitude of mIPSC events in each PC was significantly increased by 30.7% (44.72±3.78 Control vs. 58.46±6.07 pA SCF, two-tailed paired t-test p=0.0125, n= 14 cells). The potentiating effect of SCF on mIPSC amplitude was also detected by cumulative probability analysis of all individual mIPSC events (mean 45.31±0.587 vs 59.98±0.834 pA, 32.37% increase, Kolmogorov-Smirnov (KS) test p<0.0001, n=4516 Control vs 4646 SCF). Analysis of the average mIPSC kinetics per PC cell before and after SCF revealed that while rise time was unaffected, the decay was significantly decreased (Rise: 1.11±0.092 vs 1.10±0.086, two-tailed paired t-test p=0.862, n=14 pairs; Decay: 9.36±0.78 vs 6.90±0.62, two-tailed paired t-test p=0.0080, n=14 pairs.) These findings suggested that proximal GABAergic synapses undergo relative potentiation by the activation of the Kit receptor. Consistent with this interpretation, analysis of only those mIPSC events under 0.5 ms revealed a 33% increase in amplitude (Control 45.16±0.727 vs SCF 60.34±1.101 pA, KS test p<0.0001, n=2774 vs 2661). In contrast, among mIPSC events with rise times over 3 ms, SCF did not produce a significant difference in mean amplitude (Control=47.67±2.173 vs SCF 48.44±2.334, KS test p=0.0666, n=366 vs 377). Even when this threshold was expanded to include mIPSC with rise times of 2ms or greater, SCF still did not have a significant effect on mean mIPSC amplitude (Control=44.75±1.596 vs SCF 51.6±2.18, KS test p=0.1624, n=553 vs 534). Collectively, these results suggested that SCF preferentially potentiates proximal basket-cell type MLI1 mediated inhibitory output to PCs (13, 14).

We performed paired MLI1-PC recordings, choosing MLIs in the lower third of the molecular layer whose stimulation produced at least a 10% reduction in the spontaneous firing rate of PCs (6). For these MLI1-PC pairs, we measured mean PC firing rate before and after MLI1 stimulation, before vs after SCF. We analyzed this data by Two-way Repeated Measures, not assuming sphericity, with an alpha P=0.05; we determined that MLI Stimulation (P=0.0038) and SCF (P=0.0003) were significant sources of variation, while interaction of these sources was not significant (P=0.696). By Uncorrected Fisher’s LSD test, we determined that MLI stimulation before SCF (during the baseline condition) produced a significant suppression of PC firing (93.76±12.91 vs 77.29±11.26, P=0.0028), and MLI stimulation produced further suppression of PC firing after SCF application (86.67±11.77 vs 69.14±11.67 Hz, P=0.0074), 7 cell pairs. These results suggest that Kit activation increases the MLI1 mediated suppression of PC firing rate from a given amount of MLI1 stimulation, independently of effects on MLI firing per se.

We therefore determined if the application of SCF potentiated the magnitude of MLI1 evoked inhibitory currents in PCs. We injected stepped currents into MLI1s spanning 50 to 400 pA, while measuring evoked IPSC (eIPSC) in connected PCs, before and after SCF application. We analyzed sIPSC amplitude averaged from three sweeps per current step using a two-way repeated measures (per-cell) design, without assuming sphericity, with an alpha of P=0.05. We found that Current (p=0.0393) and SCF (p=0.0384), but not their interaction (p=0.2995) were significant sources of variation, using 8 MLI1-PC connected pairs. This suggests that SCF acts to increase the effectiveness of MLI1 mediated PC inhibition.

## Discussion

It has been long established that that the receptor tyrosine kinase Kit, and Kit Ligand/Stem Cell Factor (SCF) are expressed in discrete populations of synaptically coupled neurons across the brain(8, 9, 15). Our exploration of the ability of SCF to acutely tune inhibition in the cerebellum emerged from two lines of previously existing research. In one, others demonstrated that SCF can modulate excitatory synaptic communication, plasticity, and or learning in the hippocampus(7, 16, 17). In the second, we recently demonstrated through conditional knockout strategies that Kit Ligand/Kit in the developing cerebellum are essential for MLI mediated inhibition of PCs(3, 8, 15). We therefore determined if acute pharmacological manipulation of Kit signaling axis may regulate synaptic function in adult mice.

In the hippocampus, Kit Ligand is expressed in dentate gyrus granule neurons, while Kit receptor is expressed post-synaptically in CA pyramidal neurons(8). Compared to controls, mice (Sl/SL^d^ strain) carrying one null Kit Ligand allele and one allele which can only produce soluble Kit Ligand (SCF) had deficits in spatial learning: while starting at an equivalent performance, they had a longer latency after multiple days of training to find a submerged but not a visible platform and had no preference for the trained quadrant on the probe trial. In this study, the Sl/SL^d^ mice also showed lower mossy fiber-CA3 responses at baseline, but intact LTP(17). A subsequent study in control vs *Ws*/*Ws* rats carrying a 12-bp deletion in the kinase domain of Kit showed similar defects in the probe trial of a Morris maze task and defects in synaptic potentiation (16). In addition to these germline mutant studies, follow up work demonstrated that pharmacological manipulation of SCF/Kit signaling also impaired long-term changes in hippocampal plasticity (7)

In the cerebellar cortex, the biology of Kit signaling appears fundamentally different. In the hippocampus, presynaptic SCF acts on the postsynaptic CA Kit to influence glutamatergic signaling, whereas in the cerebellum, SCF is in the postsynaptic PC while Kit is in the presynaptic MLIs, both cell types GABAergic (8, 15). We recently demonstrated that conditional knockout of Kit in the embryonic hindbrain lineage that gives rise to cerebellar cortex GABAergic interneurons results in a specific loss of inhibition to PCs, apparently sparing glutamatergic input to PCs and GABAergic input to MLIs. In the same study, we found that postnatal knockout of Kit Ligand from PCs also reduces GABAergic input to PCs (3). This work implicated that Kit signaling was important for connectivity. However, unlike within certain cortical circuits, the cerebellar PC Kit Ligand and MLI Kit expression pattern is retained throughout life, which suggested a role beyond synapse development or maintenance. Drawing inspiration from the initial but limited hippocampal literature, we determined if Kit activation could acutely modulate inhibition of PCs.

We demonstrated that SCF produces an acute potentiation of inhibitory currents in PCs which requires the Kit receptor. As MLI mediated inhibition of PCs regulates PC firing rate and variability, we confirmed that SCF also produces a durable suppression of the rate and enhancement of the variability of PC firing. The nomenclature of MLIs has long involved their differential localization and morphology, with distal stellate MLIs providing inhibition to PC dendrites, and proximal basket cell MLIs providing both chemical inhibition to PC somas, and ephaptic modulation of PC output at the pinceaux formation about the PC axon initial segment. Recently, it has been further demonstrated that MLIs are segregable into at least two categories based upon their transcripts, intrinsic properties, and their connectivity. MLI1s include both morphologically stellate and basket cell types, have lower input resistance, gap junction mediated spikelets, and synapse upon other MLI1s to promote synchrony and upon PCs to mediate inhibition. In contrast, MLI12s are predominantly of a stellate morphology, have a higher input resistance, do not demonstrate gap junction mediated spikelets, and function to inhibit MLI1s to dis-inhibit PCs (11, 12, 18)We observed a suppression of PC excitability by SCF, suggesting that MLI1 output, and not MLI2 output, is facilitated by Kit activation, despite both cell types, and Purkinje Cell Layer interneurons, having Kit transcripts (12, 15, 19, 20). Supporting this interpretation, SCF did not enhance the firing rate of MLI2s nor suppress the firing rate of MLI1s, suggesting that MLI2 mediated inhibition of MLIs was not enhanced by SCF application.

The effect of Kit activation therefore seems to involve biasing MLI output to favor MLI1 mediated inhibition of PCs over MLI2 mediated PC disinhibition. Despite the presynaptic enrichment of Kit, the net effect of SCF upon inhibition of PCs seems to be primarily postsynaptic. Neither the firing rate of MLI1s, nor the frequency of mIPSC events in PCs was enhanced by SCF. Instead, the amplitude of mIPSC events was robustly increased by SCF (∼32%), with an accompanying decrease in the population decay time. The MLI1 subtypes include both distal stellate cells and proximal basket cells. The observation of a population decreased decay time suggested that the balance between distal MLI1-stellate and proximal MLI1-basket type inputs shifted to favor the latter upon SCF administration. We therefore segregated individual mIPSC events by rapid rise vs slow rises and found that only rapid-rise mIPSC events were potentiated in amplitude by SCF. We confirmed that MLI1-basket cell evoked currents in PCs were enhanced by SCF, and that SCF increased the ability of these MLI1s to further suppress PC firing, though we do not rule out the possibility of an effect (such as suppression) on MLI1 stellate cell inhibition of PCs.

Purkinje cells express both the KL1 isoform, which yields soluble SCF, and the KL2 isoform, which is membrane bound, of Kit Ligand (15). In other tiss(21)ues, these isoforms can exert different effects(22). This work with exogenous SCF suggests that increasing the relative expression or processing of KL1/SFC may favor MLI1-basket cell inhibition. Future knockout and replacement studies may inform differential roles for the KL1 and KL2 isoforms within PCs, for example in synapse modulation vs maintenance, in stellate vs basket cell type MLI1 mediated input, or in MLI2 vs MLI1 connectivity. The subcellular distribution of KL1 and KL2 isoforms within PCs is not currently known, nor are the endogenous mechanisms which may favor differential splicing, or proteolytic liberation of SCF from PCs, although such pathways have been described in other tissues (23–25). Currently, we do not know the mechanism through which PC mIPSC amplitude is increased by SCF. The presynaptic MLI Kit receptor activation may increase the quantal content of GABA in MLI1 axon terminals, or Kit receptor activation may mediate postsynaptic effects on the abundance, localization, or biophysical properties of ionotropic GABA receptors. The Kit receptor has non-catalytic roles that are not yet well understood in other tissues, and so SCF mediated dimerization of Kit may increase clustering of GABA receptors in a transsynaptic fashion to increase mIPSC amplitude . Alternatively, the activation of presynaptic Kit receptor kinase cascades may elicit effects on PC GABA_A_ receptor biology through intermediate anterograde molecules. The potential interactome of the activated Kit receptor is large, as Kit effector cascades include Ras/Raf, PI3K/Akt/mTOR, PLCγ, and Src family kinases.

Diverse neuropeptides regulate the development and function of the cerebellum, including peptides/proteins beyond SCF that play a role in synaptic modulation of PCs(26). Brain Derived Neurotrophic Factor (BDNF) enhances the inhibition of PCs; BDNF like SCF also potentiates mIPSC amplitude within PCs, though this effect seems to require postsynaptic PC TrkB receptors(27). Like SCF/Kit Ligand, BDNF also plays a role in the development of MLIs(28, 29). BDNF also modulates the development and function of climbing fibers and the plasticity of parallel fibers (paired pulse facilitation) (30). Unpublished work of others suggests that Kit signaling may also relate to the proper development or function of PC excitatory input (31). One of the first hormones ever described with canonical roles in digestion(32), the peptide secretin, is also expressed by PCs, while the G-protein coupled secretin receptor is expressed in PCs and MLIs(33). Like SCF and BDNF, secretin signaling appears to be important for the proper development of the cerebellum, mediating the survival and morphogenesis of nascent PCs and granule cells(34). We, and others, have shown that in the postnatal period secretin potentiates inhibition of PCs, (35, 36) through presynaptic mechanisms that likely involve adenyl cyclase modulation of MLI voltage gated potassium channels and AMPA receptors(37). Endogenous cerebellar secretin functions in part to modulate the acquisition of cerebellar dependent learning, including eye blink conditioning in rats(38); systemic secretin was also found in a small study to modulate eye blink conditioning in individuals with schizophrenia (39). The endogenous release of secretin from PCs is stimulated by depolarization and voltage-gated sodium and calcium channels(37). Whether PC depolarization enhances SCF liberation is not yet known.

PC depolarization is well established as a mechanism to drive multiple forms of modulation of inhibition, including transient depolarization induced suppression of inhibition (DSI), a longer depolarization induced potentiation of inhibition (DPI), and rebound potentiation (RP). While DSI is primarily mediated by PC derived retrograde cannabinoids acting upon MLI CB1 receptors, DPI involves CF derived glutamate acting at MLI NMDA receptors, both exerting a presynaptic effect. The mechanism of RP however is postsynaptic, wherein PC calcium flux enhances the function of GABA_A_ receptors through calmodulin-dependent kinase II (40–49) Since MLI derived PC inhibition is a critical gate for PC plasticity and cerebellar learning and dependent behaviors (50, 51), there must be variety of mechanisms that operate over different spatiotemporal domains and cell types to modulate inhibition. We therefore are eager to determine whether activity-induced retrograde SCF modulation of MLI1 mediated inhibition is a native mechanism for gating PC plasticity and cerebellar learning.

**Figure 1:**
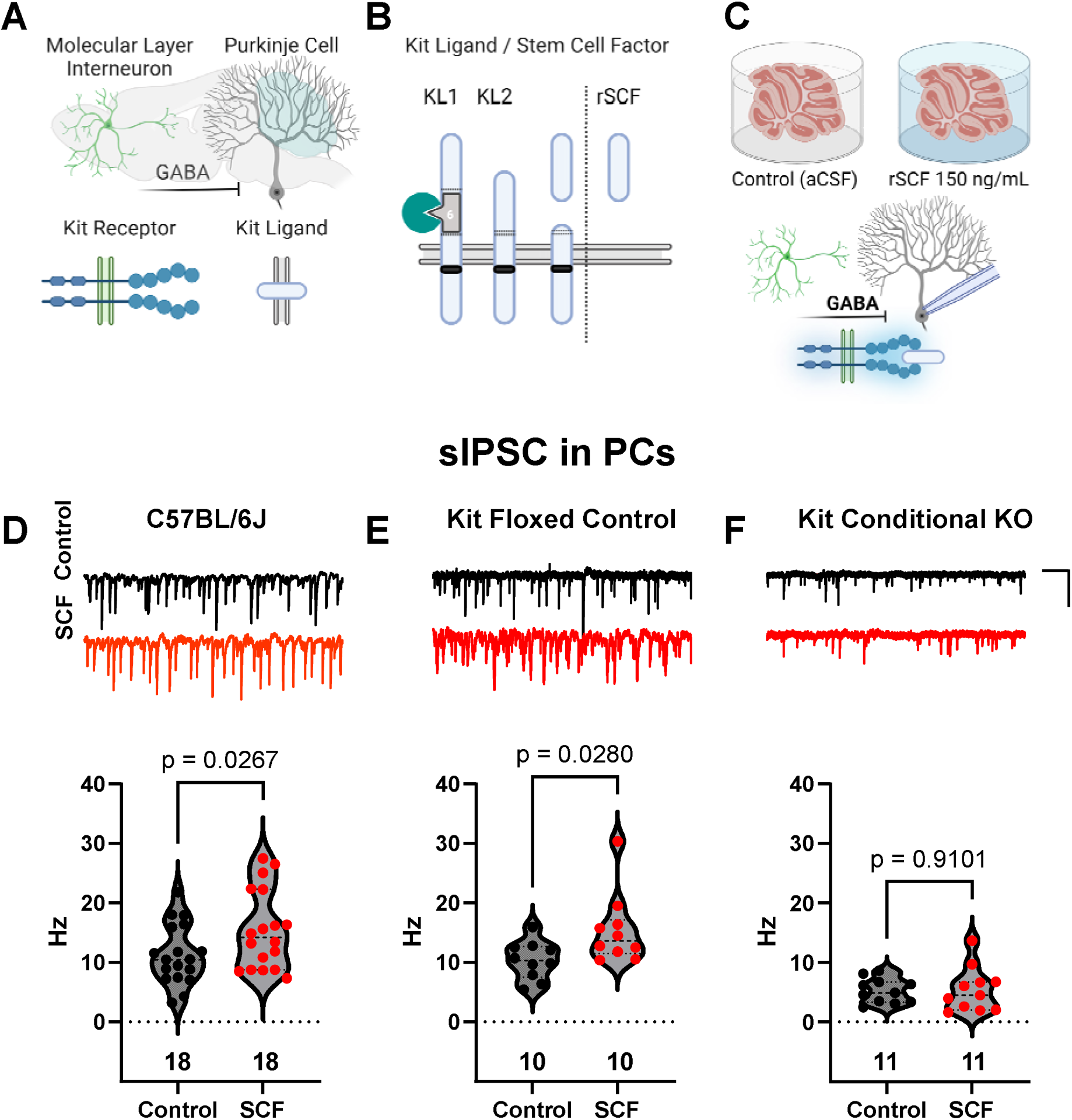
Stem Cell Factor potentiates inhibition of Purkinje Cells. **A)** Cerebellar Purkinje Cells (PCs) express the trans-membrane protein Kit Ligand, while GABAergic molecular layer interneurons (MLI) of the cerebellar cortex express the receptor tyrosine kinase, Kit. **B)** Kit Ligand encodes at least two isoforms varying in their inclusion of Exon 6, which encodes a site that facilitates the proteolytic liberation of soluble active Kit Ligand, also known as Stem Cell Factor (SCF), the recombinant form of which is an effective Kit activator. **C)** We determined if activating Kit would modulate inhibition of PCs. Acutely generated slices of mouse cerebellum were incubated in artificial cerebrospinal fluid (ACSF) alone for 30 minutes as a Control, or in ACSF supplemented with recombinant mouse SCF at 150 ng/mL for 30 minutes. From these slices, we then recorded spontaneous inhibitory postsynaptic currents in PCs. **D-F)** Representative sIPSC traces and quantification. **D)** In slices from C57BL/6J mice, SCF produced a significant increase in sIPSC frequency, and **E)** a similar effect was observed in mice on a similar genetic background which were homozygous for conditional Kit knockout (Kit Floxed) allele. However, in **F)** mice with cerebellar Kit knockout (Kit KO, homozygous for the Kit Floxed allele and Pax2 Cre positive), SCF had no effect on sIPSC frequency in PCs. **D-F)** Scale bar is 0.5 seconds by 125 pA. Statistical tests are two-tailed t-tests of the mean event frequency in each PC, n = number of PCs per condition.

**Figure 2:**
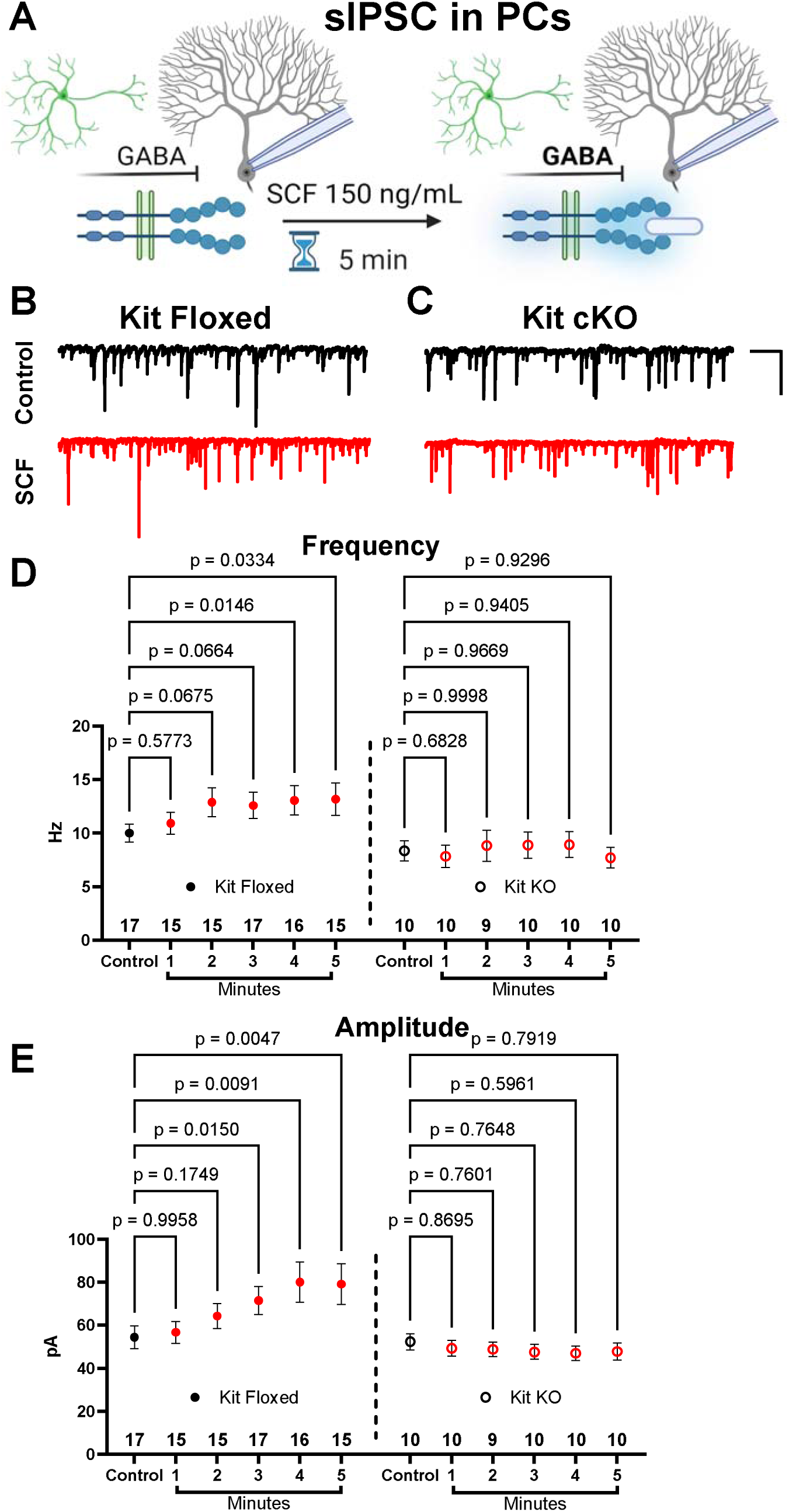
Stem Cell Factor acutely increases inhibition of Purkinje Cells. **A)** To determine the time course over which SCF potentiates inhibition, sIPSCs in PCs were analyzed in 1-minute blocks before (Control), and during a 5-minute application of 150 ng/mL SCF in both **B)** Kit Floxed and in **C)** Kit KO (Pax2 Cre mediated) animals, representative traces. **D)** From 2 through 5 minutes after the start of SCF application, the frequency of sIPSC events in PCs was significantly increased in acute cerebellar slices from Kit Floxed animals, but not in slices from Kit KO animals. **E)** Similarly, from minutes 3 through 5, the application of SCF produced a significant increase in the amplitude of sIPSC events in PC from Kit Floxed but not Kit KO animals. Scale bar is is 0.5 seconds by 125 pA. Statistical tests are mixed effects analysis of the per-PC mean sIPSC event frequency or amplitude, with Holm-Sidak multiple-comparisons test, n = number of PCs per time point per condition per genotype.

**Figure 3:**
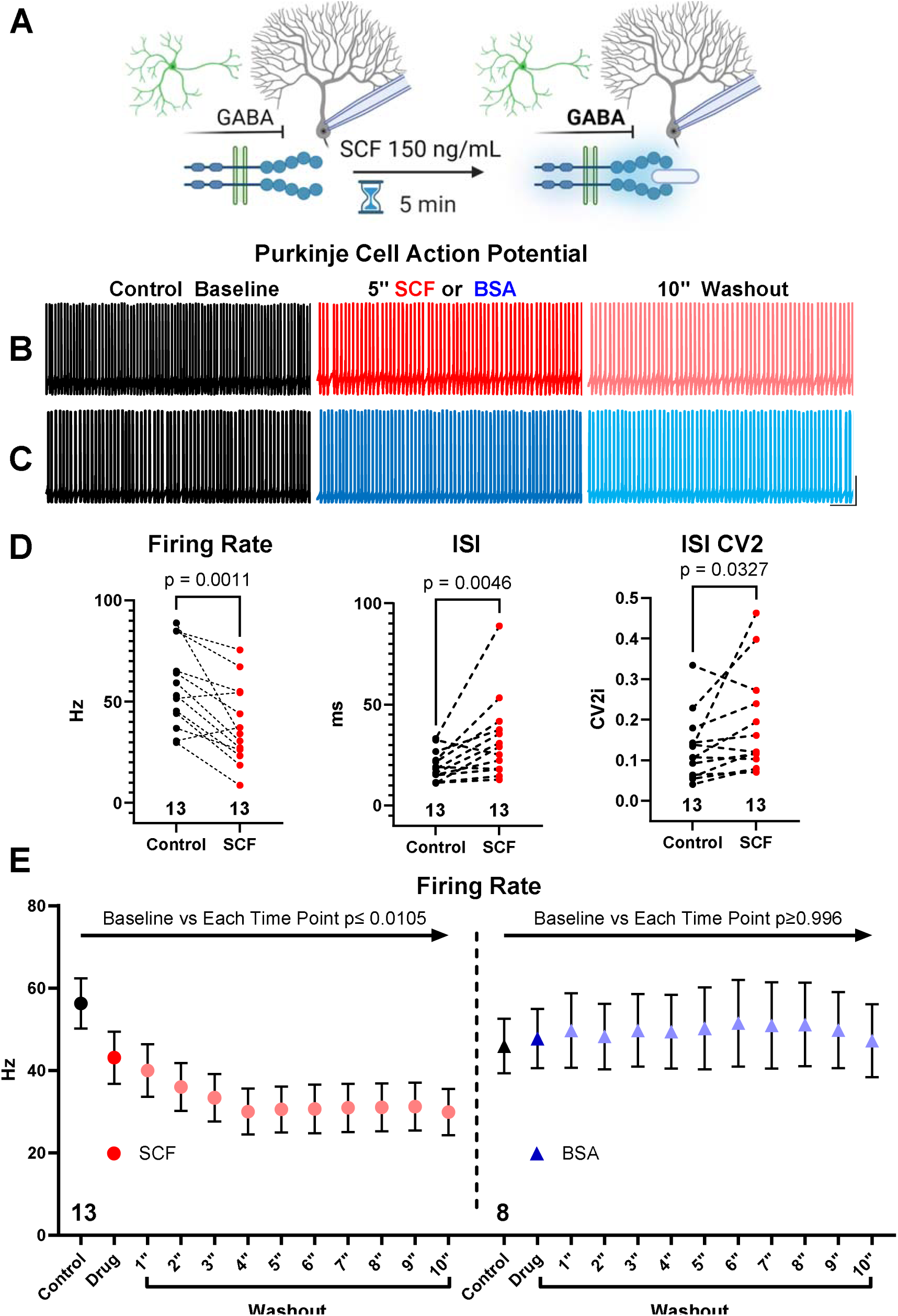
SCF induces durable suppression of PC firing. **A)** The spontaneous firing rate of PCs was determined for 1-minute periods: before (Control), after 5 minutes of 150 ng/mL SCF application, and during a 10-minute washout. **B)** Example traces of spontaneous PC action potentials during baseline Control, after 5-minute incubation in SCF or carrier solution containing bovine serum albumin (BSA), and at the end of a 10-minute Washout period. **D)** The mean firing rate of PCs was significantly suppressed by SCF application, which was accompanied by a significant increase in both the Inter-Spike Interval (ISI), as well as in the variability of the ISI (ISI CV2). **E)** The 5-minute application of SCF produces a suppression of PC firing rate that does not washout over a 10-minute period. The 5-minute application of BSA carrier does not suppress PC firing rate at any time point. Scale bar in **B)** is 125 mS by 20 mV. Graphs in **D)** reflect the per-PC means for each metric, statistical significance was determined by two-tailed t-test, except for CV2i, which was analyzed by Wilcoxon matched-pairs signed rank test. Charts for **E)** reflect separate but similar experiments, each analyzed by 2-way repeated-measures ANOVA with Holm-Sidak Multiple Comparisons test. For all statistical tests, n refers to the number of PCs per condition.

**Figure 4:**
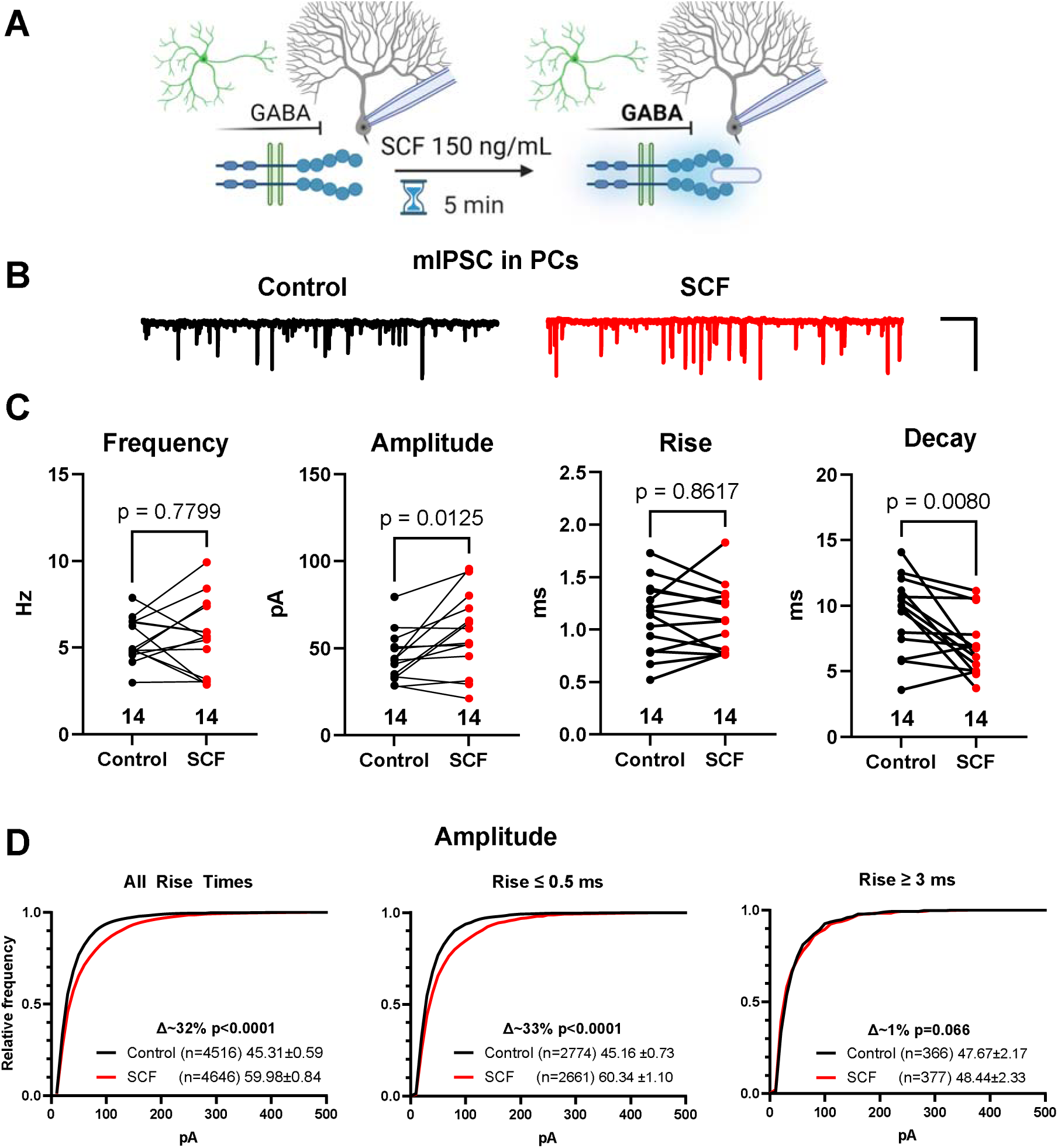
SCF potentiates proximal GABA-A synapse upon PCs. **A)** Miniature IPSCs (mIPSC) were recorded from PCs before and 5 minutes after adding SCF (150 ng/mL). **B)** Example mIPSC trace from a representative PC cell before (Control) and after SCF. **C)** The per-PC mean frequency of mIPSC events was not significantly changed by SCF. However, the average mIPSC amplitude was significantly increased by SCF, without accompanying average differences in the rise or decay times. **D)** Cumulative histograms of the proportion of individual mIPSC event amplitudes for All Rise Times, proximal basket-cell like synapses with Rise times less than 0.5 mS, and for putative stellate-cell synapses with slow rise times over 3 ms). The average amplitude of All Rise Time events, and for those with Rise <0.5 ms was significantly and similarly increased by SCF by 32-33%. However, Slow-Rise time events were not significantly increased by SCF (∼1% difference). Scale bar for **B)** is 0.5 seconds by 125 pA. Statistical analyses for **C)** are by paired two-tailed t-test where n refers to the number of PC from which mean values were derived. Statistical analyses for **D)** are by Kolmogorov–Smirnov test of n individual mIPSC event amplitudes.

**Figure 5:**
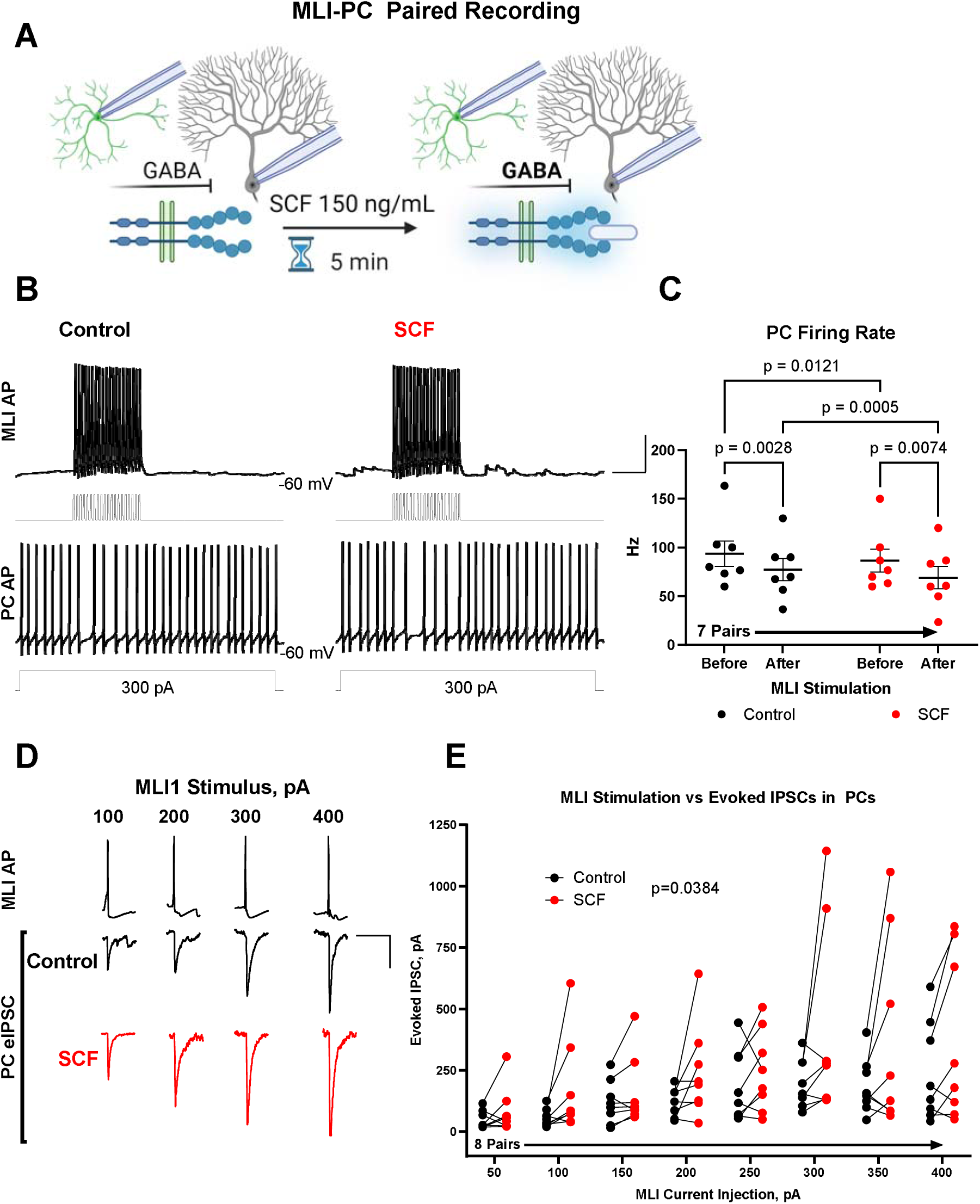
SCF potentiates MLI1 mediated inhibition of PCs. **A)** Paired recordings were made between PCs and MLI1s in the lower third of the molecular layer, connected pairs were validated by the ability of MLI1 stimulation to produce at least a 10% suppression in PC firing rate. MLI1-PC pairs were evaluated in separate experiments for either PC spiking (APs) or for evoked IPSCs, before and after SCF (5-minute, 150 ng/mL). **B)** Example traces from a connected MLI1-PC pair before and after SCF. The suppression of current-injection stimulated PC spiking was monitored before and during current-injection evoked MLI1 spiking, both before (Control) and 5 minutes after SCF application. **C)** MLI1 stimulation significantly decreased PC firing rate, and MLI1 stimulation produced a further decrease in PC firing rate after the application of SCF. **D)** Example traces of evoked AP from an MLI1 and of evoked IPSCs from a connected PC across selected MLI1 stimulus steps. **E)** The application of SCF increased the amplitude of MLI1 evoked currents recorded in PCs. Scale bar in **B)** is 50 milliseconds by 20 mV, in **E)** the scale bar is 50 milliseconds by 25mV/100pA. Statistical analysis in **C)** is repeated measures two-way ANOVA with Uncorrected Fishers Least Significant Difference; statistical analysis in **E)** is repeated-measures two-way ANOVA with Holm-Sidak multiple comparisons test; for **C) and E)** n refers to the number of MLI1-PC connected pairs evaluated before and after SCF.

---

**Figure 2—Figure supplement 1: SCF increased inhibition of PCs is mediated by GABA-A receptors.** A) sIPSC were recorded in PCs under pharmacological blockade of glutamatergic receptors (NBQX and D-AP5), glycinergic receptors (strychnine), and metabotropic GABA-B receptors (CGP-52432). The subsequent addition of SCF increased the frequency of sIPSC events, while the addition of picrotoxin nearly abolished them. Scale bar is 0.5 seconds by 125pA. Graphed data are mean sIPSC frequency per PC, analyzed by repeated-measures one-way ANOVA, Holm-Sidak multiple comparisons test, from 10 PCs across all 3 conditions.

**Figure 3—Figure supplement 1: SCF does not change MLI firing rate.** We recorded intrinsic properties and spontaneous action potential (AP) firing rate from MLIs before and after SCF (5 minutes, 150 ng/mL). **A)** An example recording of spontaneous MLI APs before and after SCF, with an input resistance of 328 MΩ. **B)** Scatter plot of MLI baseline spontaneous AP mean firing rate vs input resistance, with an arbitrary dotted line at 500 MΩ. **C)** Mean MLI AP firing rate did not significantly change after the application of SCF for either population of MLI. Scale bar is 0.5 seconds by 20 mV. Statistical significance was determined by two-tailed t-test, n refers to the number of MLIs per condition.

## Notes

### Competing Interest Statement

The authors have declared no competing interest.

