## Supplementary figures and images for "Modulation of Purkinje Cell Inhibition by Stem Cell Factor"

### Figure 2 Supplement 1

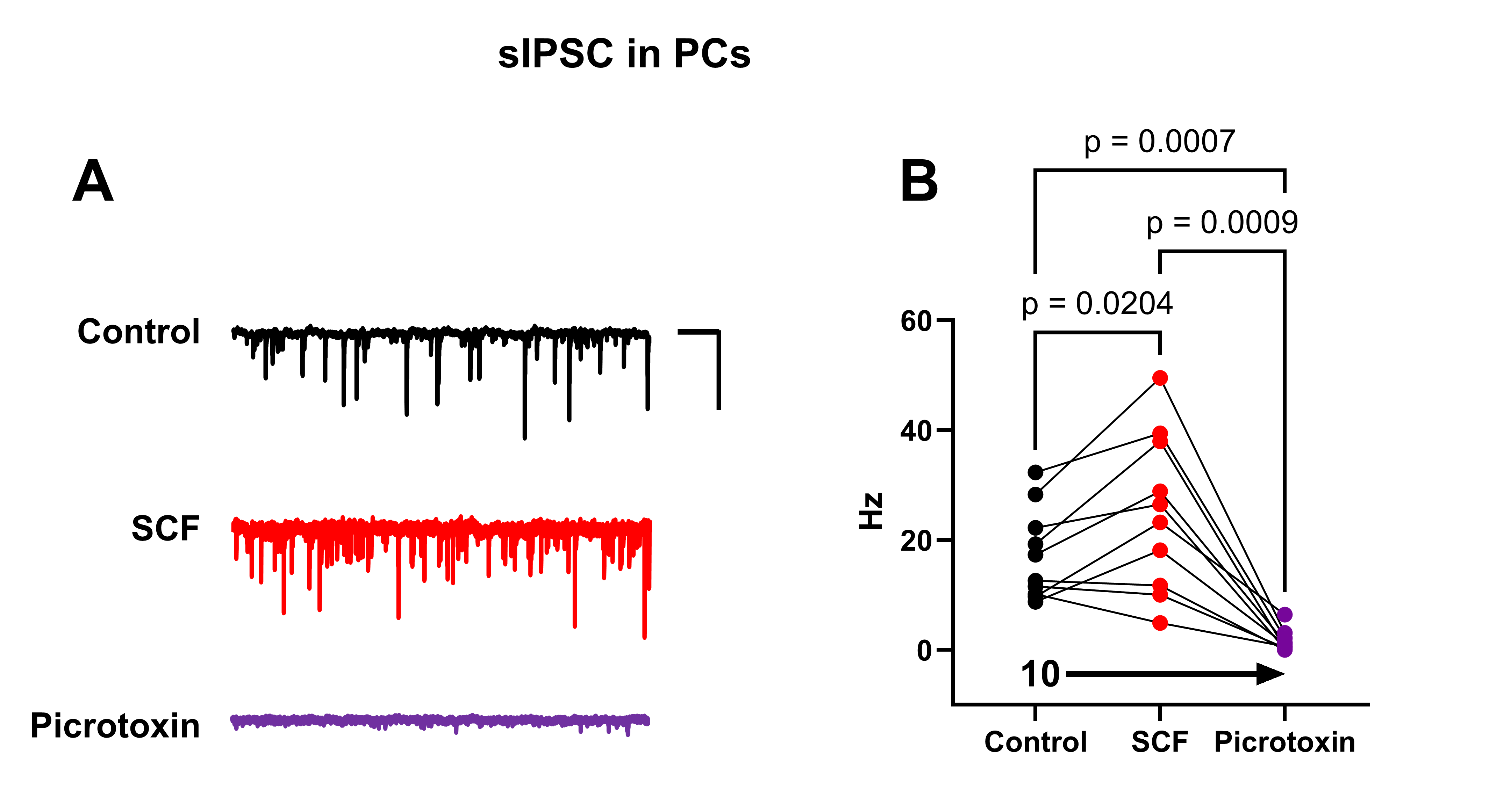

### Figure 3 Supplement 1

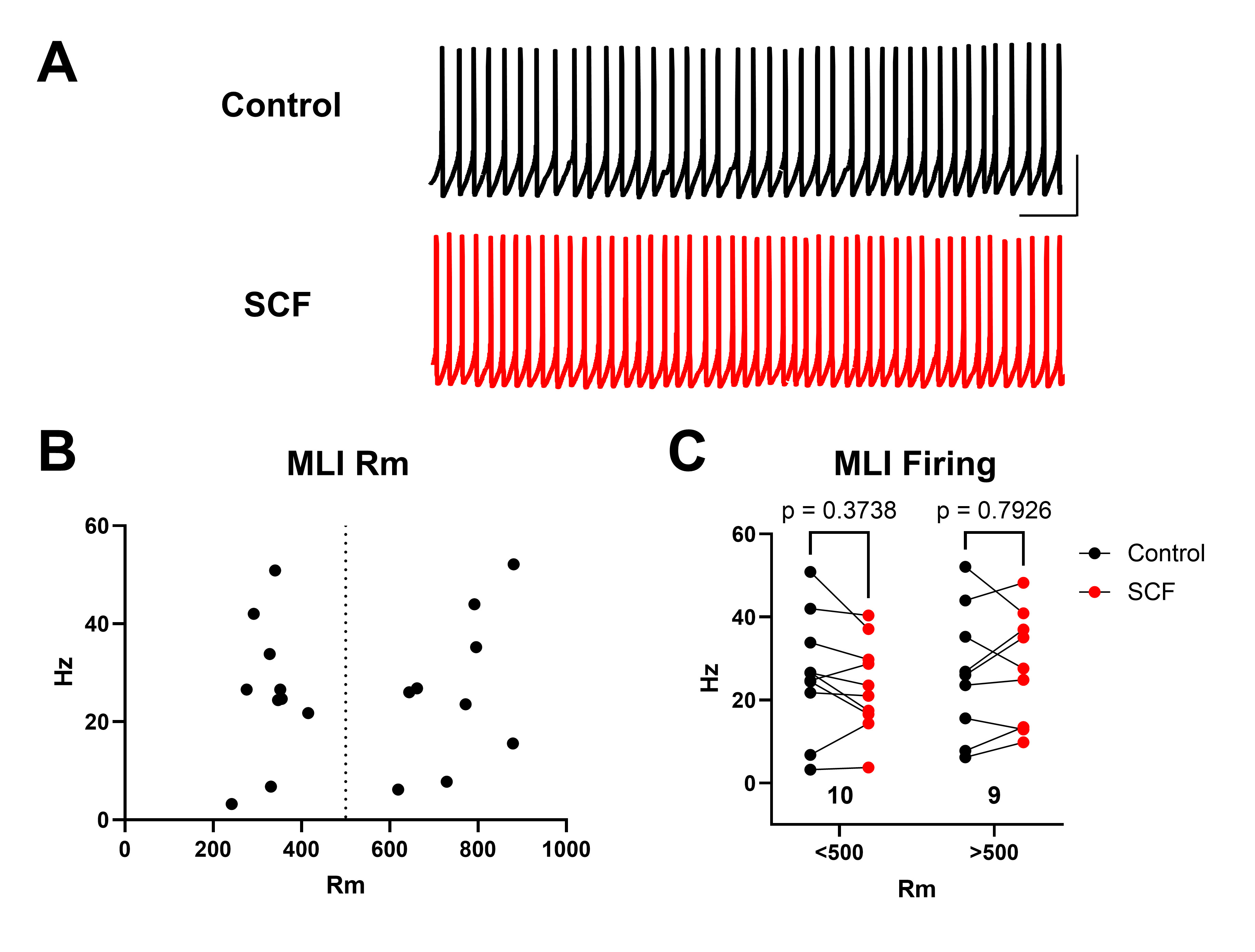
